# RTN3 inhibits RIGI-I-mediated antiviral responses by impairing TRIM25-mediated K63-linked polyubiquitination

**DOI:** 10.1101/2020.12.01.407320

**Authors:** Ziwei Yang, Jun Wang, Bailin He, Xiaolin Zhang, Xiaojuan Li, Ersheng Kuang

## Abstract

Upon viral RNA recognition, the RIG-I signalosome continuously generates IFNs and cytokines, leading to neutrophil recruitment and inflammation. Thus, attenuation of excessive immune and inflammatory responses is crucial to restore immune homeostasis and prevent unwarranted damage, yet few proresolving mediators have been identified. In the present study, we demonstrated that RTN3 is strongly upregulated during RNA viral infection and acts as an inflammation-resolving regulator. Increased RTN3 aggregates on the endoplasmic reticulum and interacts with both TRIM25 and RIG-I, subsequently impairing K63-linked polyubiquitination and resulting in both IRF3 and NF-κB inhibition. Rtn3 overexpression in mice causes an obvious inflammation resolving phenomenon when challenged with VSV, Rtn3-overexpressing mice display significantly decreased neutrophil numbers and inflammatory cell infiltration, which is accompanied by reduced tissue edema in the liver and thinner alveolar interstitium. Taken together, our findings identify RTN3 as a conserved proresolving mediator of immune and inflammatory responses and provide insights into the negative feedback that maintains immune and inflammatory homeostasis.

**Summary:** RTN3 is upregulated upon RNA viral infection due to inflammation and ER stress, in turn suppresses antiviral responses by impairing TRIM25-mediated RIG-I K63-linked polyubiquitination and consequently decreases neutrophil populations and inflammatory infiltration, representing a novel mechanism of negative inflammatory resolution.

## Introduction

The innate immune responses can be triggered by pathogen-associated molecular patterns (PAMPs) and then serve as the first line of defense against invading pathogens ^1^. Upon viral infection, pattern recognition receptors (PRRs) detect viral RNA, DNA and other viral products and subsequently mediate the activation of downstream signaling pathways ^2 3^. Both RIG-I and MDA5, two crucial members of the RIG-I-like receptor (RLR) family, sense cytoplasmic viral RNA and activate antiviral responses. Both RIG-I and MDA5 share the same domain architecture, comprising two N-terminal caspase activation recruitment domains (CARDs), a central DEAD box helicase/ATPase domain and a C-terminal regulatory domain (CTD) ^3^. MDA5 has been reported to recognize longer dsRNA molecules (>2 kb), while RIG-I prefers shorter (< 1-2 kb) or 5’ triphosphate-containing dsRNA ^4 5^. Unlike MDA5, which is conformationally unaltered, upon recognition of dsRNA by the CTD domain, RIG-I unfolds and releases its CARDs from the inward folding state formed by its helicase domain, thereby transforming into an activated state ^6,7^. The CARD domain then recruits MAVS and activates the downstream signaling cascade to induce the expression of type I interferons, IFN stimulated genes (ISGs), inflammatory cytokines and/or chemokines ^7^.

Considering that RIG-I-mediated innate antiviral responses are important, drastic and rapid upon viral RNA recognition, multiple mechanisms have evolved to precisely regulate RIG-I-mediated antiviral signaling to maintain the balance between immunity and tolerance and prevent severe or even fatal unnecessary damage, which may be caused by excessive immune and inflammatory responses ^8^. Some modifications of RIG-I reduce its stability. For example, LRRC25 targets RIG-I for ISG15-associated autophagy degradation ^9^, while the E3 ubiquitin ligases RNF125, STUB1, c-Cbl and CHIP induce the K48-linked polyubiquitination of RIG-I to promote its proteasome-dependent degradation ^10 11^. Alternatively, the posttranslational modification of RIG-I upon K63-linked polyubiquitination has been well shown to regulate RIG-I activation. TRIM25 and Riplet induce this modification in RIG-I ^12 13^, while the deubiquitination enzyme USP3 targets the RIG-I CARDs to cleave its K63-linked polyubiquitin ^14^. In addition, CKII and PKCα/β phosphorylate the RIG-I CTD and CARDs to negatively regulate its activation ^15 16^.

Although the acute immune response and inflammation defend against viral infection and eliminate virus-induced damage, excessive, unrestricted responses can lead to excessive tissue injury and even organ failure. Several diseases have been shown to be related to chronic inflammation or autoimmunity, such as asthma, arthritis, periodontal disease and neurodegenerative disorders ^17 18 19 20^. Inflammation resolution has been shown to be an essential active process ^21^, and proresolving mediators regulate inflammation resolution through multiple strategies, including the inhibition of signal transduction, the regulation of cytokines and chemokines, the cessation of leukocyte infiltration and the clearance of apoptotic cells ^19 22^.

Viral infections also trigger different ER stresses due to the excessive synthesis and accumulation of viral proteins in the ER lumen or through signal hijacking ^23^. In response to ER stress, two transcription factors, CHOP and ATF6, are activated, and the translational regulator eIF2α is phosphorylated and subsequently reprograms cellular transcription and translation ^24 25^. Although reticulon family members have shown to comprise a highly conserved housekeeping protein family in multiple species ^26^, one reticulon member, RTN3, is transcriptionally upregulated under ER stress by CHOP or ATF6 ^27^.

In our previous unpublished experiments, we observed that RTN3 is dramatically upregulated upon RNA viral infection when using VSV as a stimulus. However, the mechanism and function of RTN3 upregulation during viral infection is unclear, raising the strong possibility that RTN3 is involved in antiviral and inflammatory responses. In the present study, we showed that RTN3 acts as a proresolving mediator during RNA viral infections and suppresses RIG-I-mediated immune and inflammatory responses. We reveal that RTN3 exerts an inhibitory effect on RIG-I signalosome activation by impairing TRIM25-mediated RIG-I K63-linked polyubiquitination. Our study may contribute to a better understanding of a novel mechanism for inflammation resolution and the reconstitution of tissue homeostasis during viral infections.

## Results

### RTN3 is upregulated and self-aggregates upon RNA viral infection

In our previous study, we occasionally observed increased RTN3 expression upon vesicular stomatitis virus (VSV) infection, leading us to investigate why RNA viral infection increases RTN3 expression. Wild-type (WT) HEK293T cells were infected with enhanced GFP-tagged vesicular stomatitis virus (VSV-eGFP) or treated with the dsRNA synthetic analog poly(I:C) at different time points. We observed that RTN3 protein levels were dramatically increased during VSV-eGFP infection (Fig. 1 A, top) or under poly(I:C) stimulation (Fig. 1 A, bottom), with both conditions showing the same pattern of *RTN3* mRNA upregulation (Fig. 1 B, 1 C, top) accompanied by increased *TNF*-α mRNA levels that were used to indicate the effectiveness of VSV-eGFP or poly(I:C) treatment towards inflammatory induction in stimulated cells (Fig. 1 B, 1 C, bottom). Interestingly, we observed that TNF-α could upregulate RTN3 expression at both the protein (Fig. 1 D) and mRNA levels (Fig. 1 E). These results suggest that RTN3 levels are increased during RNA viral infection.

**Figure 1.**
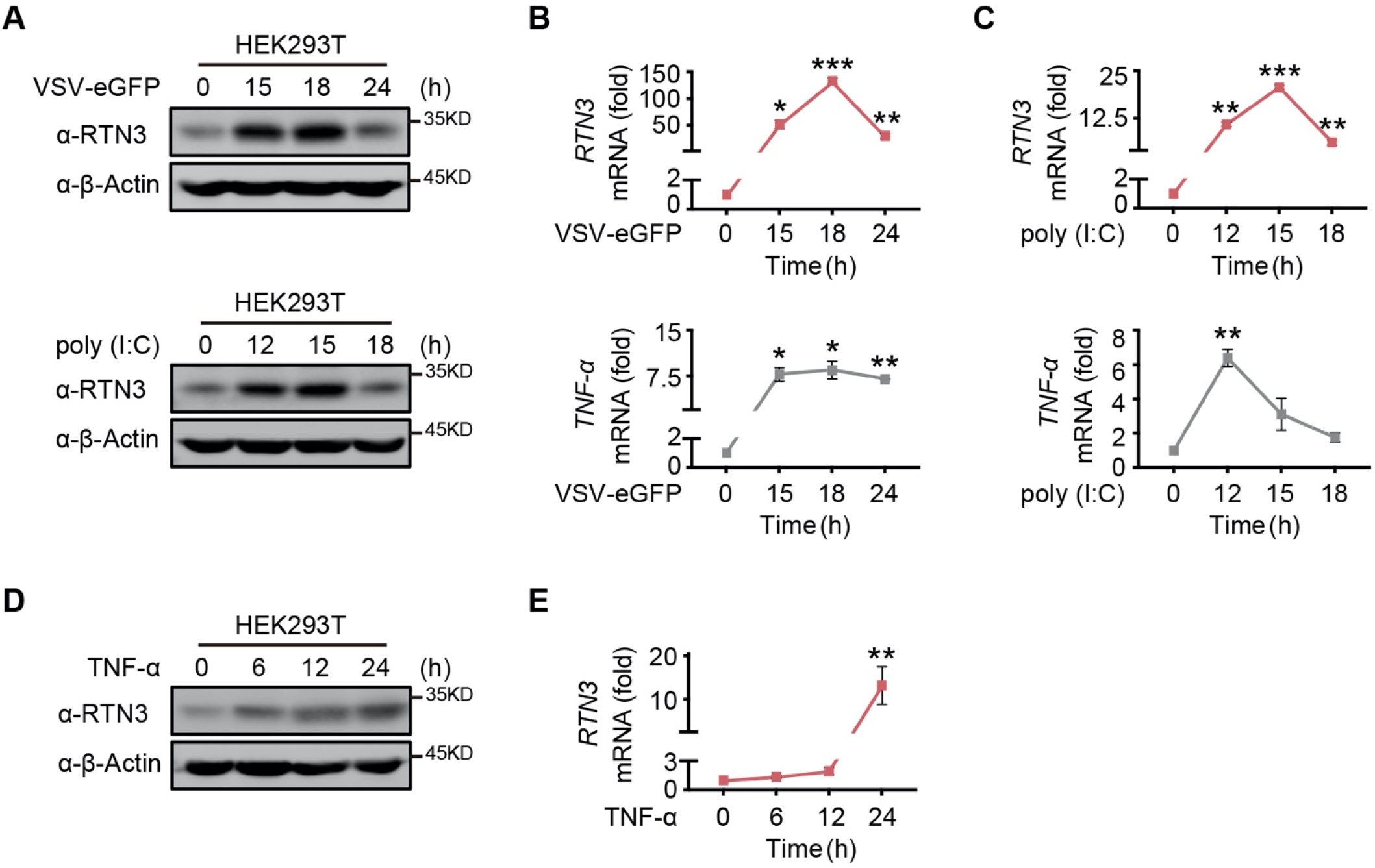
RTN3 is upregulated and self-aggregates upon RNA viral infection. A. Immunoblot analysis of HEK293T cells infected with VSV-eGFP (MOI = 1) or treated with poly(I:C) (5 μg/ml) at the indicated timepoints. B-C. mRNA levels of *RTN3* and *TNF*-α in the same samples shown in (A) were detected by real-time PCR. B. VSV-eGFP-infected group, C. poly(I:C)-treated group. D. Immunoblot analysis of HEK293T cells treated with TNF-α (10 ng/ml) at the indicated timepoints. E. mRNA levels of RTN3 in the same samples shown in (D) were detected by RT-PCR. In (A, D), the data are representative of three independent experiments. In (B, C, E), the data are shown as the mean values ± SD (n = 3). *, p < 0.0332; **, p < 0.0021; ***, p < 0.0002; and ****, p < 0.0001 by Sidak’s multiple comparisons test.

Since RTN3 is conserved and ubiquitously expressed in various tissues, we further assessed the pattern of RTN3 upregulation in multiple cell lines, including A549 and THP-1 cells. As expected, the protein (Fig. S1 A, S1 B) and mRNA (Fig. S1 C) levels of RTN3 were both significantly increased by poly(I:C) stimulation or VSV-eGFP infection in A549 cells, with a similar phenomenon observed in THP-1 cells (Fig S1 D, S1 E, S1 F). Notably, confocal microscopy analysis of HeLa cells transfected with mCherry-tagged RTN3 followed by challenge with poly(I:C) showed that RTN3 was localized on the endoplasmic reticulum (ER), while under poly(I:C) stimulation, RTN3 proteins converged on the ER and formed many aggregated bodies of varying sizes (Fig. S1 G). Taken together, these data suggest that RTN3 expression is induced by RNA viral infection and its observed upregulation and self-aggregation indicate that RTN3 may be involved in regulating innate immune and inflammatory responses.

### RTN3 overexpression suppresses antiviral immune responses

To investigate the function of RTN3 in regulating innate immune responses, we performed luciferase assays using ISRE-luc, IFNβ-Luc and NF-κB-Luc reporters and observed that RTN3 markedly inhibited all activities induced by RIG-I overexpression (Fig. 2 A). In addition, poly(I:C)- or Sendai virus (SeV)-stimulated ISRE-luc activities were both restrained by RTN3 overexpression in a dose-dependent manner (Fig. 2 B). However, RTN3 overexpression had a slightly negative effect on MDA5-induced ISRE-luc activities (Fig. S2 A) and barely inhibited TLR3-induced ISRE-luc activities (Fig. S2 B). These data reveal that RTN3 primarily suppresses RIG-I-mediated antiviral immune responses.

**Figure 2.**
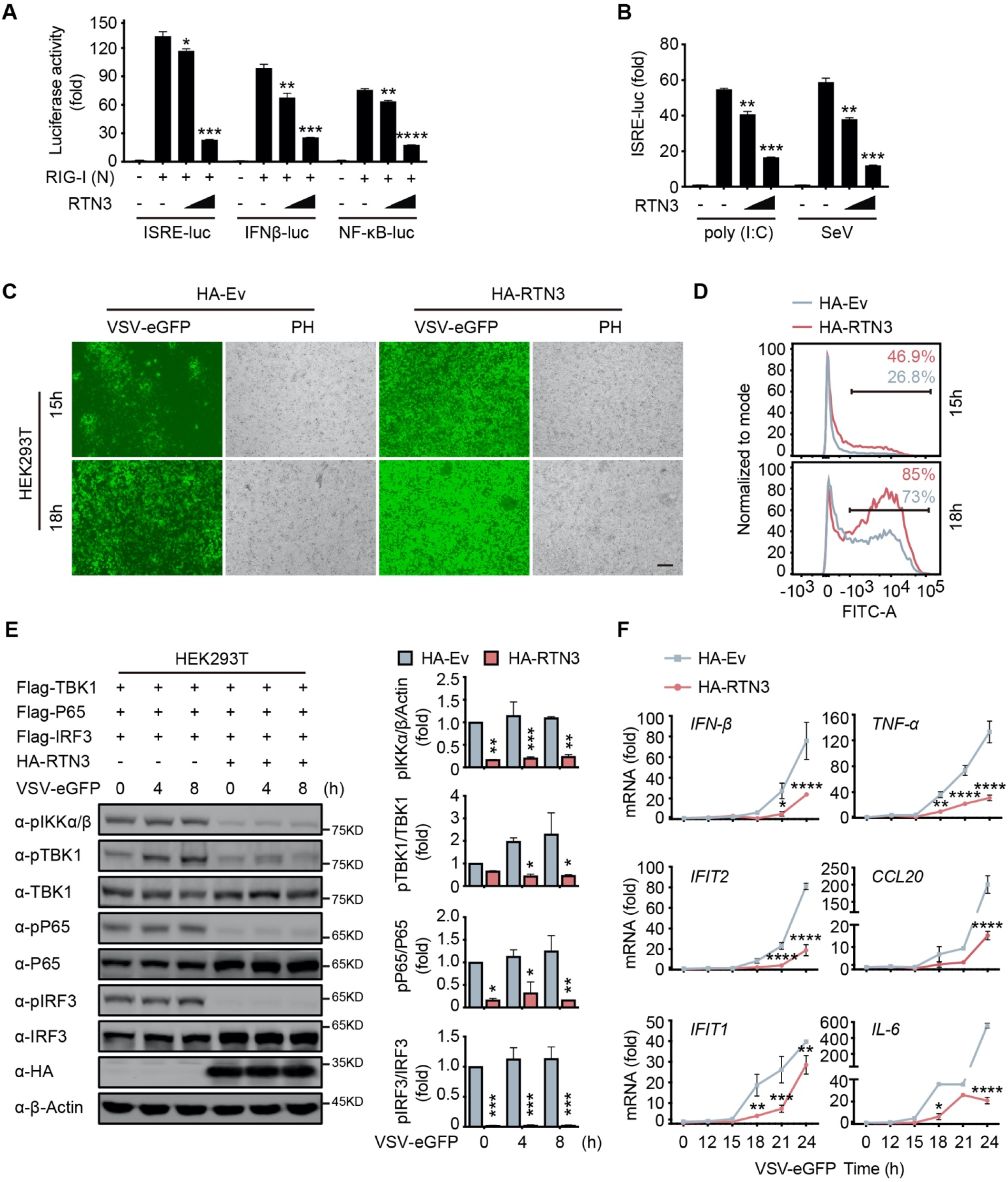
RTN3 overexpression suppresses antiviral immune responses. A. Luciferase assays of HEK293T cells transfected with an ISRE luciferase reporter (ISRE-Luc), an IFNB luciferase reporter (IFNβ-Luc) or an NF-κB luciferase reporter (NF-κB-Luc) together with an HA-tagged empty vector (HA-Ev, no wedge) or increasing amounts (wedge) of plasmid encoding RTN3 (HA-RTN3) followed by transfection with a plasmid encoding the RIG-I CARD domain [RIG-I (N)] to activate the pathway. Cell lysates were collected 24 h post transfection, the same as that described in the following experiments, if no additional annotation was performed. B. Luciferase activity of HEK293T cells transfected with ISRE-Luc, together with HA-Ev or increasing amounts of plasmid HA-RTN3, followed by treatment with poly(I:C) (5 μg/ml) or SeV (MOI = 0.1) for 16 or 24 h. C. Fluorescence and phase contrast (PH) analyses of HEK293T cells transfected with HA-Ev or HA-RTN3, followed by infection with VSV-eGFP (MOI = 0.05) at the indicated timepoints. Scale bar, 100 μm. D. The percentage of eGFP-positive cells in the same samples shown in (C) was analyzed by flow cytometry. E. Immunoblot analysis of HEK293T cells transfected with HA-Ev or HA-RTN3 together with plasmids encoding TBK1, NF-κB p65 and IRF3 (Flag-TBK1, Flag-p65 and Flag-IRF3) followed by infection with VSV-eGFP (MOI = 1) at the indicated timepoints (left). Quantitative comparison of the indicated protein levels analyzed by gray intensity scanning of blots (right). F. RT-PCR analysis of *IFN-*β, *IFIT2, IFIT1, TNF-*α, *IL-6* and *CCL20* mRNA levels in HEK293T cells transfected with HA-Ev or HA-RTN3 followed by infection with VSV-eGFP (MOI = 1) at the indicated timepoints. In (E), the data are representative of three independent experiments. In (A, B, F), the data are shown as the mean values ± SD (n = 3). *, p < 0.0332; **, p < 0.0021; ***, p < 0.0002; and ****, p < 0.0001 by Sidak’s multiple comparisons test.

To further demonstrate the attenuation of antiviral activities by RTN3, HEK293T cells were transfected with an RTN3-encoding plasmid and subsequently infected with VSV-eGFP. Fluorescence microscopy and flow cytometry results showed that RTN3 overexpression rendered the cells highly susceptible to viral infection compared to the empty vector-transfected cells, which displayed a less sensitive phenotype (Fig. 2 C, 2 D). A consistent result was also observed in long-term puromycin-selected, stable HA-tagged RTN3 vs. HA-tagged empty vector-expressing THP-1 cells (THP-1^HA-RTN3^ vs. THP-1^HA-Ev^) (Fig. S2 D), which excluded the possibility that ectopic RTN3 expression may transiently overload the endoplasmic reticulum (ER) and impair the translation of antiviral proteins.

To further assess the role of RTN3 in attenuating antiviral activities, we transfected HEK293T cells with the minimum amount of TBK1-, NF-κB p65-, and IRF3-encoding plasmids as primers together with RTN3 or empty vector and followed by infection with VSV-eGFP. The phosphorylation of IKKα/β, TBK1, p65 and IRF3 was dramatically decreased in RTN3-overexpressing cells compared to that observed in the control groups (Fig. 2 E, left). Quantitative analysis by gray intensity scanning of immunoblot bands further demonstrated the significant decreases in the phosphorylation of the two kinases and two transcription factors (Fig. 2 E, right).

Accordingly, we investigated whether RTN3 suppresses type I IFN- and interferon-stimulated gene (ISG) expression. RT-PCR results showed that RTN3-overexpressing cells exhibited a deficiency in *IFN-*β, *IFIT2, IFIT1, TNF-*α and the proinflammatory cytokines *CCL-20* and *IL-6* during VSV infection (Fig. 2 F). Similarly, the mRNA levels of *IFN-*β, *IFIT2* and *IFIT1* were downregulated to a great extent in THP-1^HA-RTN3^ cells compared to THP-1^HA-Ev^ cells after VSV infection (Fig S2 E). Because the amounts of the proinflammatory cytokines CCL-20 and IL-6 were both markedly decreased by RTN3 overexpression under dsRNA stimulation [VSV or poly(I:C)], we assessed whether any other inflammatory cytokines or related genes were altered by RTN3 overexpression. To this end, we performed RT-PCR array analysis and observed that many inflammatory factors and genes were downregulated during RTN3 overexpression (Fig. S2 F). These results suggest that RTN3 overexpression preferentially suppresses RIG-I-mediated antiviral immune and inflammatory responses.

### RTN3 interacts with TRIM25 or RIG-I

Next, we investigated the potential mechanism for the RTN3-mediated inhibition of antiviral responses. We noted that RTN3 may function as a tripartite motif-containing protein 25 (TRIM25)-binding protein through mass spectrometry analysis ^28^, and TRIM25 is a crucial E3 ligase for the K63-linked polyubiquitination modification of RIG-I to promote its activation upon RNA viral infection ^12^. To elucidate the relationship between RTN3 and TRIM25 or RIG-I, HEK293T cells were transfected with GFP-tagged RTN3 and Flag-tagged TRIM25 followed by stimulation with or without VSV and then subjected to coimmunoprecipitation and immunoblot analysis. The results showed that RTN3 interacted with TRIM25 under basal conditions and was slightly strengthened upon viral infection (Fig. 3 A, Fig. S3 A). Considering that the RTN3 overexpression leads to its aggregation in the ER and may cause nonspecific interactions (such as structural enfolding), we used GFP-TRIM25 to pull down endogenous RTN3 and confirmed its interaction with TRIM25 and that this interaction was strengthened by VSV infection (Fig. 3 B). Since TRIM25 directly targets RIG-I, we speculated that RTN3 may also interact with RIG-I. Therefore, we performed coimmunoprecipitation assays and observed an interaction between TRIM25 and RIG-I (Fig. 3 C) as well as a similar VSV-promoting pattern as that detected for RTN3 interacting with TRIM25 (Fig. S3 B). Furthermore, confocal microscopy results showed the colocalization of RTN3 with RIG-I or TRIM25 on the extended ER framework (Fig. 3 D), indicating that RTN3 aggregation may provide a scaffold for TRIM25 oligomerization or the RIG-I signalosome complex. Collectively, these results suggested that RTN3 interacts with both RIG-I and TRIM25 and that this interaction could be enhanced by RNA viral infection.

**Figure 3.**
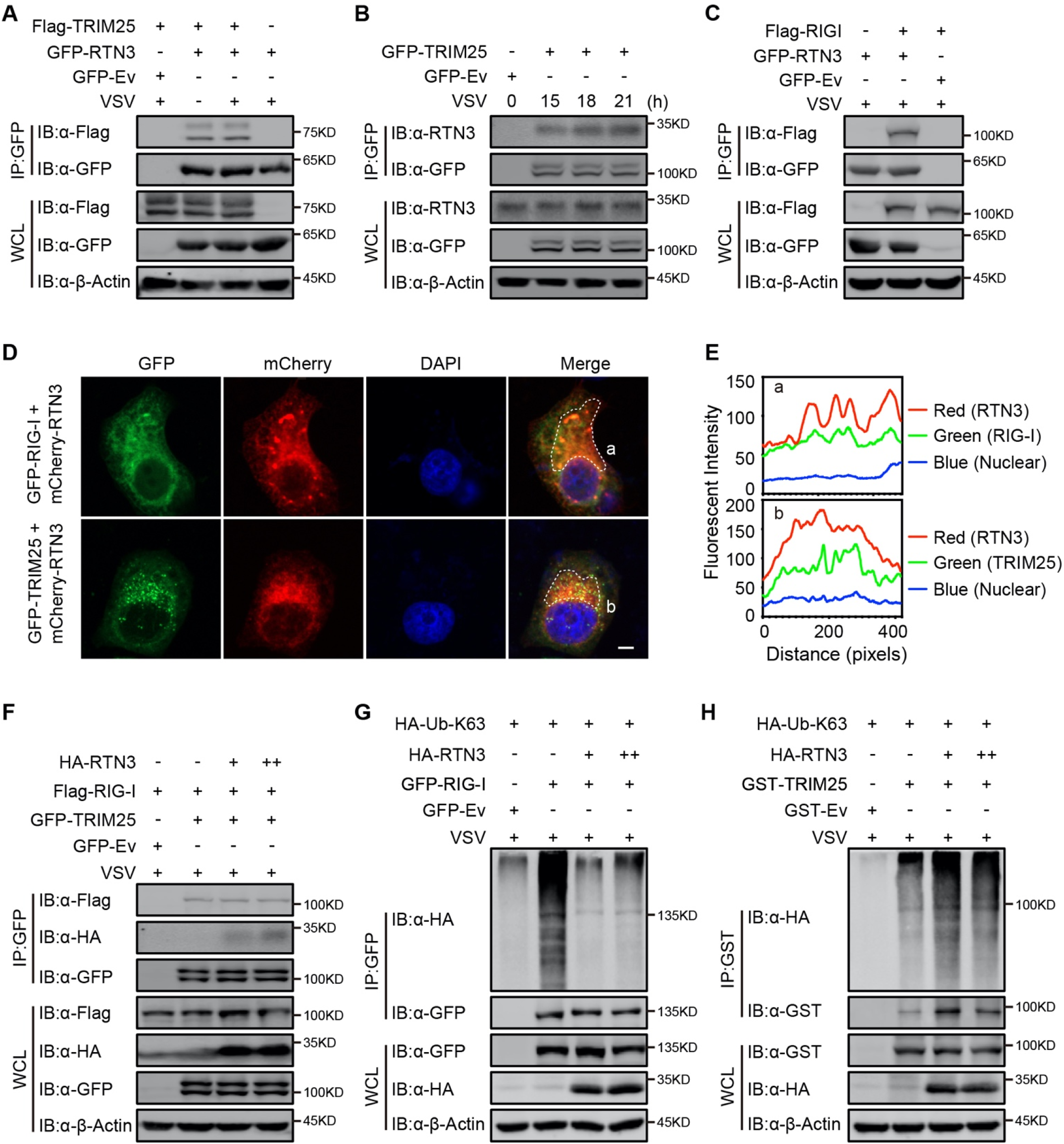
RTN3 separately interacts with TRIM25 and RIG-I and inhibits the K63-linked polyubiquitination of RIG-I. A. Coimmunoprecipitation (CoIP) and immunoblot analyses of HEK293T cells transfected with a plasmid encoding TRIM25 (Flag-TRIM25) or a Flag-tagged empty vector (Flag-Ev) together with GFP tagged RTN3 (GFP-RTN3) or a GFP-tagged empty vector (GFP-Ev) in the indicated groups for 24 h and followed by infection with VSV (MOI = 1) for 8 h. B. Immunoprecipitation (IP) and immunoblot analyses of HEK293T cells transfected with GFP-TRIM25 or GFP-Ev in the indicated groups followed by infection with VSV (MOI = 1) at the indicated timepoints. C. CoIP and immunoblot analyses of HEK293T cells transfected with a plasmid encoding RIG-I (Flag-RIG-I) or Flag-Ev together with GFP-RTN3 or GFP-Ev in the indicated groups for 24 h and followed by infection with VSV (MOI = 1) for 8 h. D. Confocal microcopy analysis of HeLa cells transfected with GFP-RIG-I and mCherry-tagged RTN3 (mCherry-RTN3) or GFP-TRIM25 with mCherry-RTN3. The dotted box (a, b) indicates the colocalization area. Scale bar, 5 μm. E. Qualitative analysis of the fluorescence intensity of the selected area in (D--a, b). F. CoIP and immunoblot analyses of HEK293T cells transfected with HA-Ev or increasing amounts of HA-RTN3 (+, ++) together with Flag-RIG-I and GFP tagged TRIM25 (GFP-TRIM25) or GFP-Ev in the indicated groups for 24 h and followed by infection with VSV (MOI = 1) for 8 h. G. CoIP and immunoblot analyses of HEK293T cells transfected with HA-Ub-K63 and HA-Ev or increasing amounts of HA-RTN3 (+, ++) together with GFP-RIG-I or GFP-Ev in the indicated groups for 24 h and followed by infection with VSV (MOI = 1) for 8 h. H. CoIP and immunoblot analyses of HEK293T cells transfected with HA-Ub-K63 and HA-Ev or increasing amounts of HA-RTN3 (+, ++) together with GST tagged TRIM25 (GST-TRIM25) or GST-Ev in the indicated groups for 24 h and followed by infection with VSV (MOI = 1) for 8 h. In (A-C) and (F-H), the data are representative of three independent experiments.

### RTN3 impairs the K63-linked polyubiquitination of RIG-I upon RNA viral infection

We next assessed how RTN3 suppresses RIG-I-mediated antiviral responses through its interaction with TRIM25. Interestingly, the protein levels of RIG-I and TRIM25 were not decreased by RTN3 overexpression in HEK293T cells (Fig. S3 C, S3 D), excluding the possibility that RTN3 may target RIG-I or TRIM25 for their degradation. We also assessed whether the interaction between RIG-I and TRIM25 was inhibited or promoted by RTN3 overexpression. To this end, HEK293T cells were cotransfected with Flag-RIG-I- and GFP-TRIM25-expressing constructs or GFP-tagged empty vector (GFP-Ev) together with increasing amounts of RTN3-expressing plasmid, which was followed by VSV infection. Unfortunately, the interaction between RIG-I and TRIM25 was completely unchanged (Fig 3 F), indicating that RTN3 negatively regulates RIG-I-mediated antiviral responses via other mechanisms.

Subsequently, we investigated the polyubiquitination of RIG-I upon VSV-eGFP infection in the absence or presence of RTN3 overexpression. As shown in Fig. 3 G, the K63-lined ubiquitination level of RIG-I was greatly decreased by RTN3 overexpression, potentially suppressing RIG-I signalosome formation and downstream signal transduction. We further investigated the ubiquitination of TRIM25 in the presence of RTN3, as the autoubiquitination (K63-linked) of TRIM25 is essential for its catalytic activity ^29^. Following the increasing expression of RTN3 in HEK293T cells, the K63-linked polyubiquitination of TRIM25 was significantly promoted (Fig. 3 H), consistent with the observed RTN3-mediated increase in the polyubiquitination of TRIM25 (Fig. S3 E). Taken together, these data indicate that RTN3 suppresses RIG-I-mediated antiviral innate immune responses by inhibiting K63-linked polyubiquitination upon viral RNA recognition, simultaneously suppressing the K63-linked ubiquitination of RIG-I and promoting the K63-linked polyubiquitination of TRIM25.

### RTN3 inhibition of antiviral responses is TRIM25-dependent

To elucidate whether TRIM25 promotes the ability of RTN3 to inhibit RIG-I-mediated immune responses, we further investigated whether TRIM25 deficiency abolishes the inhibition of RIG-I-mediated antiviral immune responses. To this end, endogenous TRIM25 expression was depleted in HEK293T cells by shRNA (HEK293T-shTRIM25), after which the phosphorylation of IKKα/β, TBK1 and p65 was detected in both HEK293T-shTRIM25 and HEK293T-shCtrl cells. Compared to the HEK293T-shCtrl cells, TRIM25 depletion counteracted the inhibition of IKKα/β, TBK1 and p65 phosphorylation induced by RTN3 overexpression when challenged with VSV-eGFP (Fig. 4 A). Similarly, RTN3 overexpression in HEK293T-shTRIM25 cells only slightly impaired RIG-I K63-linked polyubiquitination upon VSV infection (Fig. 4 B). Taken together, these data suggest that TRIM25 has a crucial role in the RTN3-mediated suppression exerted of RIG-I activation.

**Figure 4.**
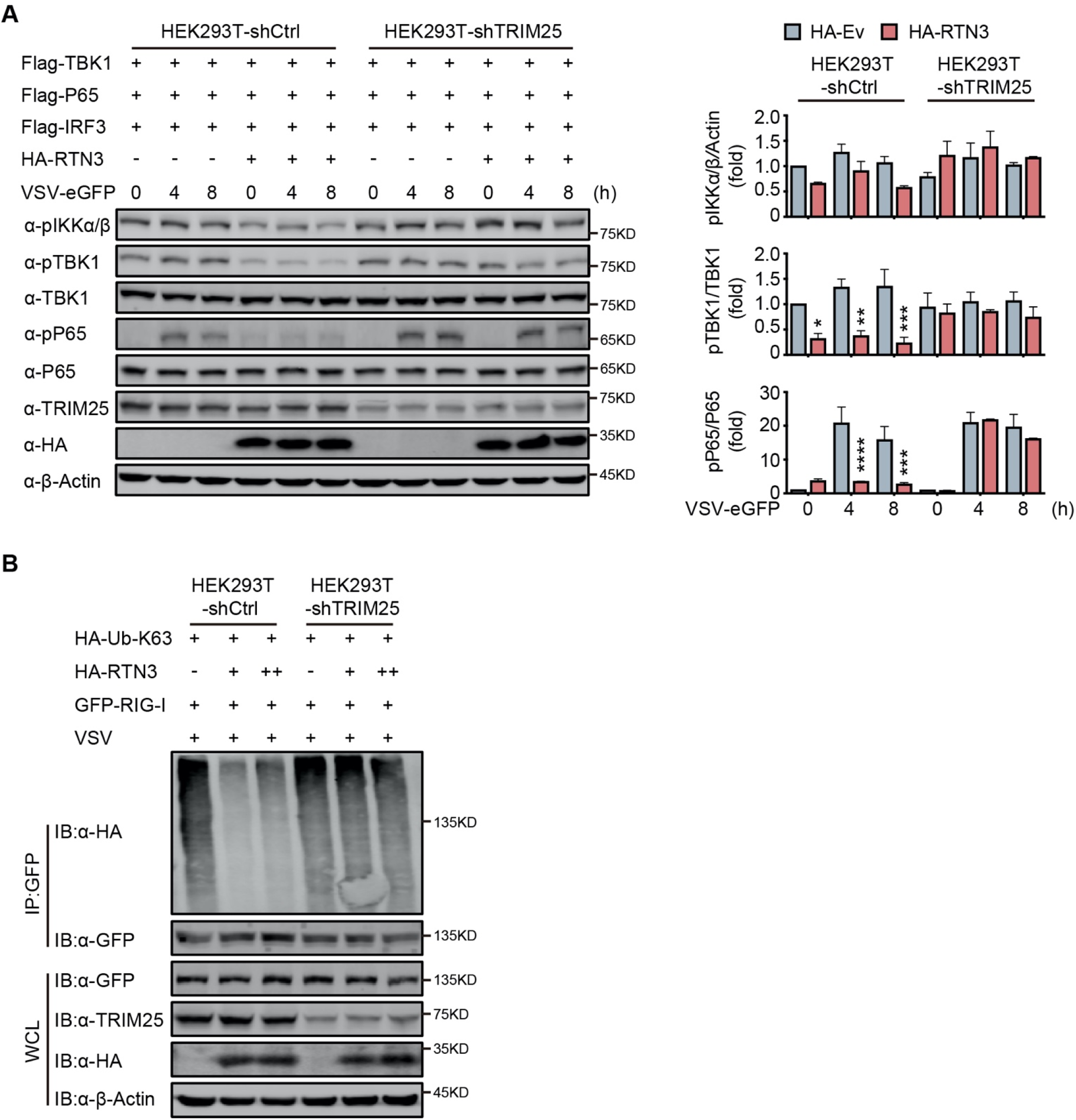
RTN3-mediated inhibition of antiviral responses is TRIM25-dependent. A. Immunoblot analysis of control HEK293T cells (HEK293T-shCtrl) and TRIM25 knockdown HEK293T cells (HEK293T-shTRIM25) transfected with HA-Ev or HA-RTN3 together with Flag-TBK1 and Flag-p65 and followed by VSV-eGFP infection (MOI = 1) at the indicated timepoints (left). Quantitative comparison of the indicated protein levels in (A) analyzed by gray intensity scanning of blots (right). B. CoIP and immunoblot analysis of HEK293T-shCtrl and HEK293T-shTRIM25 cells transfected with HA-Ub-K63 and increasing amounts of HA-RTN3 (+, ++) or HA-Ev together with GFP-RIG-I or GFP-Ev in the indicated groups for 24 h and followed by infection with VSV (MOI = 1) for 8 h. In (A, B), the data are representative of three independent experiments. In (A, left), the data are shown as the mean values ± SD (n = 6). *, p < 0.0332; **, p < 0.0021; ***, p < 0.0002; and ****, p < 0.0001 by Sidak’s multiple comparisons test.

Subsequently, we evaluated which domain of RTN3 is responsible for its inhibitory activity towards RIG-I-mediated antiviral responses. RTN3 localizes to the ER membrane and comprises a noncytoplasmic domain (NC) that inserts into the ER lumen, three transmembrane domains (TM) and two cytoplasmic domains. A series of RTN3 truncations were constructed from the C terminus based on its domain structure (Fig. S4 A) and were then assessed for their ability to suppress RIG-I-mediated immune activation through ISRE-luc and NF-κB-luc reporter assays. The TM1-3-truncated construct (T4 mutant) was unable to inhibit RIG-I (N)-induced ISRE-luc activity, while other truncated mutants maintained similar inhibitory activities as full-length RTN3 (Fig. S4 B, top). The same phenomenon was also observed in NF-κB-luc reporter assays (Fig. S4 B, bottom). These results suggest that the first TM domain is required for the inhibitory activity of RTN3. Furthermore, ubiquitination analysis demonstrated that K63-linked polyubiquitination was impaired by the RTN3 mutants T1, T2 and T3 at levels similar to wild-type RTN3, whereas T4 mutants lacked this activity (Fig. S4 C). Collectively, our data suggest that RTN3 inhibits RIG-I-mediated innate immune antiviral responses in a TRIM25-dependent manner and that transmembrane domain 1 (aa 61-92) of RTN3 is indispensable for this activity.

### RTN3 overexpression suppresses antiviral immune responses in mice

We next assessed whether RTN3 is upregulated and inhibits RIG-I-mediated innate immune antiviral responses in vivo during viral infection by treating C57BL/6 mice with PBS, poly(I:C) or VSV-eGFP separately via intravenous (I.V.) injection. Real-time PCR results showed that the mRNA levels of *Rtn3* in liver extracts were significantly higher in the VSV- and poly(I:C)-treated mice than in those of the PBS control mice (Fig. 5 A), and Rtn3 protein levels were also markedly upregulated (Fig. 5 B). These results confirm that RTN3 expression is significantly induced by RNA viral infection in mice, at least in the liver.

**Figure 5.**
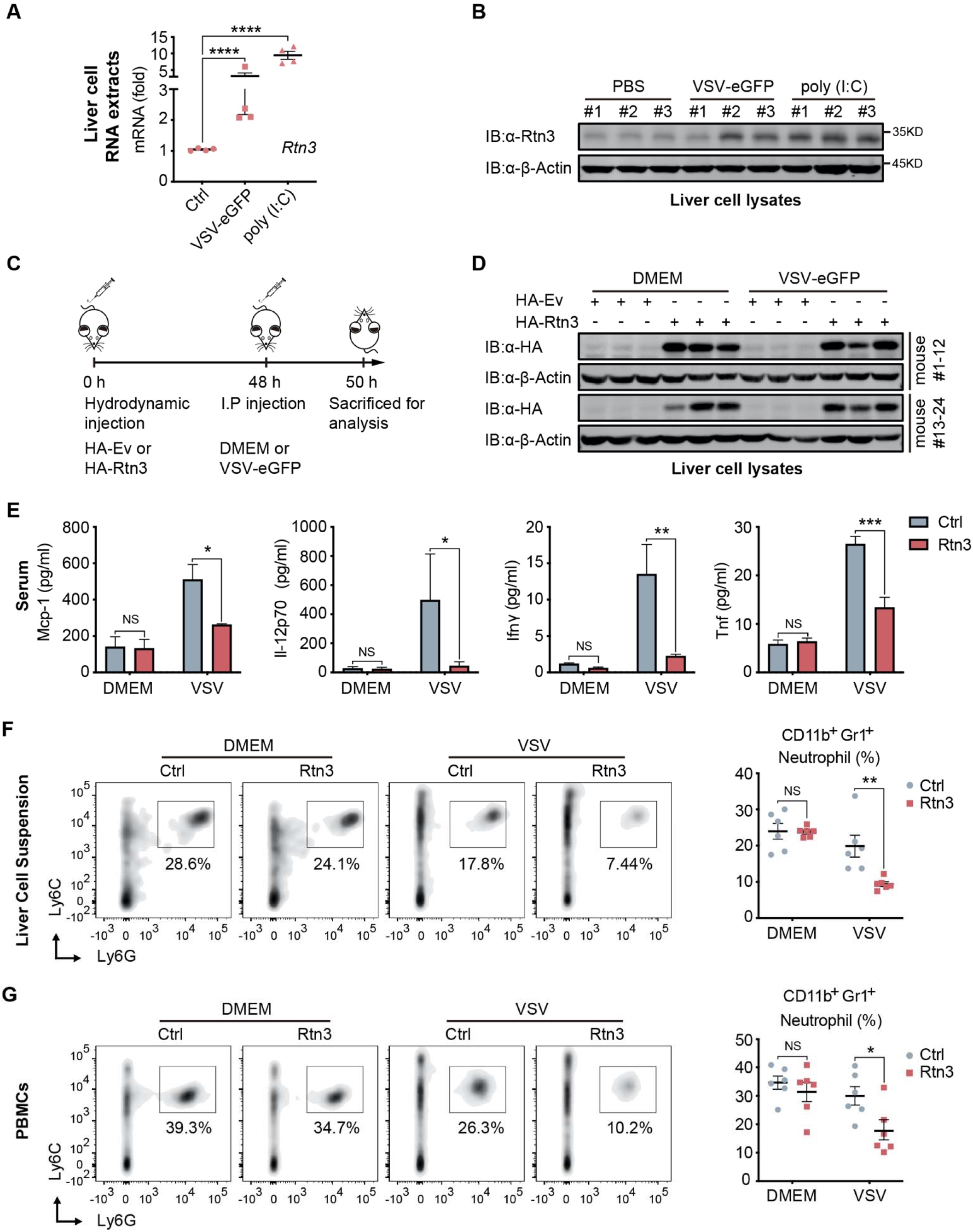
RTN3 overexpression suppresses antiviral immune responses in mice. A. mRNA levels of *Rtn3* were detected by RT-PCR in liver cells from mice treated with PBS, poly(I:C) (400 µg/mouse) or VSV-eGFP (1×10^7^ PFU/g/mouse) for 12 h. B. Immunoblot analysis of liver cell lysates of the same samples shown in (A), with “# number” indicating individual mice. C. Timeline of in vivo experiments. For plasmid hydrodynamic injection, 20 µg/mouse; for viral infection (intraperitoneal injection), 1×10^7^ PFU g^-1^/mouse VSV (VSV-eGFP) or 600 µl DMEM /mouse. D. Immunoblot analysis of liver cell lysates from the same samples shown in (C). E. Protein levels of Mcp-1, IL-12p70, Ifn-γ and Tnf-α in serum samples from the mice shown in (C) were measured by flow cytometry using a BD™ CBA Mouse Inflammation kit. F. The percentage of neutrophils in liver cell suspensions from the same mice shown in (C) was detected by flow cytometry. The density graph indicates the cell population groups for each mouse (left), and the percentage comparison for all mice is shown in the scatterplot (right). G. The percentage of neutrophils among PBMCs from the same mice as in (C) was detected by flow cytometry. The density graph indicates the cell population grouped of each mouse (left), and the percentage comparison for all mice is shown in the scatterplot (right). In (A, E, F, G), the data are shown as the mean values ± SD (n = 4 in A, n = 6 in E-G). *, p < 0.0332; **, p < 0.0021; ***, p < 0.0002; and ****, p < 0.0001 by Sidak’s multiple comparisons test.

To further investigate whether RTN3 upregulation inhibits antiviral responses in vivo, mice were hydrodynamically injected with a plasmid encoding Rtn3 (HA-Rtn3) or the empty vector (HA-Ev) and then inoculated with VSV-eGFP via intraperitoneal (I.P.) injection to infect mice for another 12 h (Fig. 5 C). At 48 h, as expected, appropriate levels of plasmid-mediated RTN3 overexpression were detected in the livers of mice (Fig. 5 D). After the mice were sacrificed and then bled, the serum samples were incubated with precoated beads, and flow cytometry results showed that the levels of key inflammatory factors, including Mcp-1, IL-12p70, IFN-γ and TNF-α were consistently decreased in the Rtn3-overexpressing mice upon viral infection, especially IFN-γ and TNF-α, both of which were markedly downregulated (Fig. 5 E). We next analyzed the immune cell population within the liver and PBMCs through flow cytometry analysis. Cells were grouped as indicated (Fig. S5 A, S5 B), and upon VSV-eGFP infection, the population of CD11b^+^Gr1^+^ neutrophils (Fig 5 F, 5 G) dramatically decreased in the Rtn3-overexpressing mice, while other immune cell populations, including CD11^+^Ly6C^hi^F480^+^ macrophages, Ly6C^hi^F4/80^lo^ monocytes, CD45^+^CD3^+^ T cells and CD45^+^CD3^-^CD11b^+^CD11c^+^ dendritic cells, showed no significant change (Fig. S5 C). Taken together, these results indicate that upon RNA viral infection, RTN3 is upregulated and subsequently suppresses antiviral immune and inflammatory responses, decreasing the population of neutrophils in mice.

### RTN3 overexpression suppresses inflammation in mice

Based on the observed inhibitory activity of RTN3 in mouse experiments, we speculated that RTN3 may promote the resolution of inflammation in vivo. To assess this possibility, the liver and lung tissue sections of mice were stained with hematoxylin and eosin (H&E), and microcopy analysis showed that upon VSV infection, RTN3 overexpression (indicated by RTN3 IHC) alleviated inflammatory cell infiltration and tissue edema in the liver (Fig. 6 A top and middle, 6 B, 6 C), which was accompanied by a thinning of the alveolar interstitium in the lung (Fig. 6 A bottom, 6 D). These results indicated that RTN3 upregulation is induced by RNA viral infection to attenuate inflammation and protect against tissue injury caused by excessive inflammatory responses. Further experiments evaluating VSV-eGFP infection in the liver revealed that the VSV-infected cell population was increased in RTN3-overexpressing mice (Fig. S6 A, S6 B), and the MFI of the GFP signal indicated that VSV-eGFP infection also increased to a certain extent. Collectively, these results indicate that RTN3 upregulation under RNA viral infection attenuates tissue inflammation and likely facilitates the resolution of inflammation.

**Figure 6.**
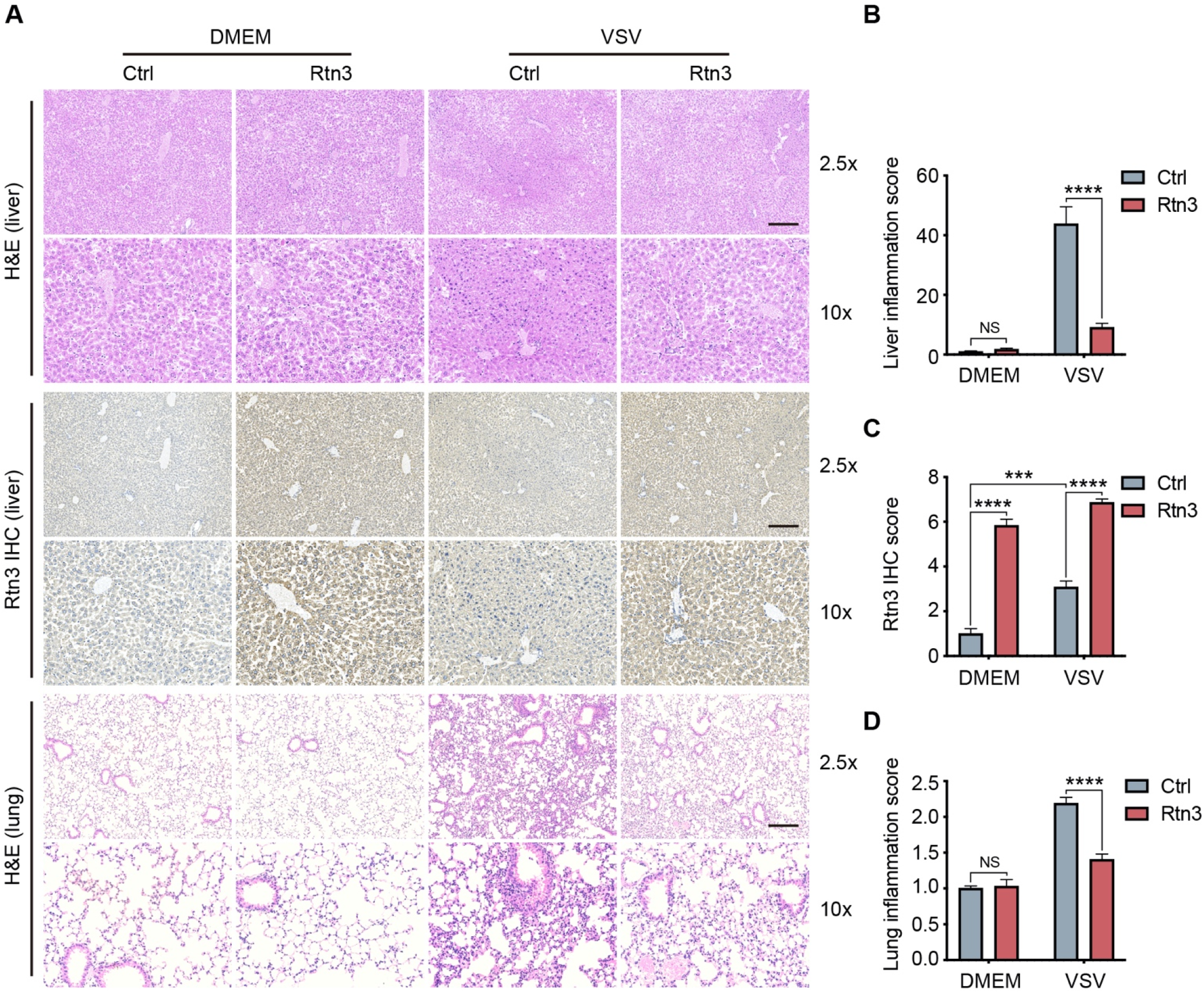
Rtn3 overexpression promotes inflammation resolution in mice. A. Microcopy analysis of liver and lung tissue sections from the same mice as in (Fig. 5 C) with hematoxylin and eosin (H&E) staining, and HA-Rtn3 exotic expression was visualized by immunohistochemical staining (IHC). B. Inflammation scores of liver tissue sections (n=4, 4, 4, 4) from the same mice shown in (A). The data were obtained and analyzed using ImageJ and visualized with GraphPad Prism 8. C. IHC score assessment of liver tissue sections as in (A, middle), data was obtained and calculated with Image J plugin “IHC Profiler” D. Inflammation scores of lung tissue sections (n=4, 4, 4, 4) from the same mice shown in (A). The data were obtained and analyzed using ImageJ and visualized with GraphPad Prism 8. In (B-D), data are shown as the mean values ± SD (n = 4). *, p < 0.0332; **, p < 0.0021; ***, p < 0.0002; ****, p < 0.0001; by Sidak’s multiple comparisons test.

## Discussion

In the present study, with respect to the conserved and ubiquitous expression and ER subcellular localization of reticulon members, we demonstrate that RTN3 is involved in the negative regulation of antiviral responses upon RNA viral infection. Our results demonstrate that RTN3 is upregulated upon infection with VSV or poly(I:C) stimulation and then self-aggregates along the ER membrane. A potential mechanism for RTN3 induction may be ER stress upon acute viral infection. As a UPR target gene, the promoter region of the *RTN3* gene harbors binding sites for the ER stress-related transcription factors ATF6 and CHOP. As a consequence, RTN3 upregulation inhibited RIG-I-mediated immune responses, the poly(I:C)-, VSV- or SeV-induced expression of type I IFNs and cytokines, and activated of the IRF3 and NF-κB pathways but had only a slight and minimal effects on MDA5- and TLR3-induced activation, respectively. We further demonstrated that RTN3 overexpression significantly attenuates the RIG-I-mediated production of proinflammatory cytokines.

In assessments of whether RIG-I is the predominant target through which RTN3 inhibits antiviral responses, our results demonstrated that RTN3 separately interacts with RIG-I and TRIM25 and that their interaction is augmented upon VSV stimulation. Furthermore, we showed that RTN3 overexpression impairs the K63-linked polyubiquitination of RIG-I in a TRIM25-dependent manner but does not interfere with RIG-I stability or interaction. Notably, the K63-linked polyubiquitination of TRIM25 was promoted while that of RIG-I was inhibited by RTN3 overexpression that led to RTN3 self-aggregation and the formation of a scaffold-like structure, indicating that RTN3 may interfere with the oligomerization of TRIM25 and its E3 ubiquitin ligase activity to promote RIG-I polyubiquitination. Furthermore, mapping analysis of truncated RTN3 constructs showed that RTN3 transmembrane domain 1 (TM1) is crucial for impairing RIG-I K63-linked polyubiquitination and inhibiting immune responses, indicating that a proper topological structure of RTN3 on the ER membrane is essential for its activity. Thus, the underlying mechanism of how RTN3 interferes with this process is worth further investigation.

Consistent with the in vitro results, in vivo experiments in mice showed that RTN3 is upregulated upon VSV infection or poly(I:C) stimulation. Importantly, our results provide evidence that RTN3 overexpression decreases the production of inflammatory factors and the neutrophil population in the liver and blood during RNA viral infection. The native neutrophil population may be primarily affected by the downregulation of cytokines and chemokines, and the circulating neutrophil population is affected secondarily. Because neutrophils have the capacity to damage tissue ^30 31^, neutrophil inhibition is crucial in the inflammatory response ^22 32^. The suppression of chemokine CXCL8 (IL-8; Fig. S2) levels by RTN3 indicates that it may be a potential mediator of inflammation resolution by indirectly restricting neutrophil recruitment. Although the decreased macrophage populations in Rtn3-ovrerexpressing mice were not significant, we did observe a decreasing trend for these populations (Fig S5 C). The histological results demonstrated that RTN3 overexpression attenuated the infiltration of lymphocytes and greatly relieved tissue inflammation and edema. Taken together, these results provide evidence that RTN3 functions to maintain tissue homeostasis and initiate the resolution of inflammation by inhibiting RIG-I-mediated antiviral responses and even terminating acute inflammation.

Therefore, we propose a working model that describes how RTN3 suppresses RIG-I-mediated antiviral innate immune responses (Fig. S7). Upon RNA viral infection, the RIG-I signalosome undergoes TRIM25-mediated activation and induces the production of IFNs, cytokines and chemokines, leading to acute immune responses and inflammation. Subsequently, RTN3 is upregulated in an ER stress- and/or inflammation-dependent manner, which is directly associated with acute viral infection. Aggregated RTN3 on the endoplasmic reticulum interacts with TRIM25 and RIG-I, inhibiting the K63-linked polyubiquitination of RIG-I via the E3 ligase activity TRIM25 to block the RIG-I signaling cascade and attenuate innate immune responses. As a result, the production of IFNs, cytokines and chemokines is attenuated, and the resolution of host inflammation is simultaneously initiated.

In summary, the results of our present study demonstrate that a conserved reticulon protein, RTN3, which is ubiquitously expressed in various tissues and organs, is upregulated upon RNA viral infection. In turn, upregulated RTN3 inhibits RIG-I signalosome activation by impairing the TRIM25-RIG-I axis. Our findings demonstrate a novel negative feedback mechanism for acute immune responses and inflammation resolution and the potential reconstitution of tissue homeostasis during viral infection.

## Materials and Methods

### Mice and mouse experiments

All animal experiments were approved by the Institutional Animal Care and Use Committee of Sun Yat-Sen University, China. Wild-type (WT) C57BL/6 mice were purchased from VITAL RIVER, Beijing, China and used for the animal experiments. For Rtn3-induction models, 8-week-old female mice received an intravenous (I.V.) injected of poly(I:C) (400 μg/mouse) or an equal volume of phosphate-buffered saline (PBS) as controls or VSV-eGFP [1 × 10^7^ plaque-forming units (PFU)/g/mouse)] for 12 h. For RTN3-overexpressing mouse models, 8-week-old female mice were hydrodynamically injected with a plasmid encoding Rtn3 (HA-Rtn3) or the empty vector (HA-Ev). Twenty micrograms of DNA diluted in 2 ml of 1× PBS was injected through the tail vein using a syringe with a 30-gauge needle. The injection was completed within 8 to 12 seconds, and 48 h post DNA injection, the mice received an intraperitoneal (I.P.) injection of VSV-eGFP [1 × 10^7^ PFU/g/mouse)] or an equal volume of DMEM for 12 h.

### Cell culture

HEK293T and HeLa cells were cultured in DMEM (Corning, 10-013-CV) supplemented with 10% fetal bovine serum (FBS) (Gibco, 10270-106), while A549 and THP-1 cells were cultured in RPMI 1640 medium (Corning, 10-040-CV) supplemented with 10% FBS. THP-1 cells stably overexpressing HA-RTN3 (THP-1^HA-RTN3^) and control THP-1 cells (THP-1^HA-Ev^) were cultured in RPMI 1640 medium supplemented with 10% FBS, while TRIM25 knockdown HEK293T cells (shTRIM25-HEK293T) were cultured in DMEM supplemented with 10% FBS. All cells were cultured in an incubator at 37°C under an atmosphere with 5% CO_2_.

### Antibodies and reagents

The following antibodies were used in this study: anti-RTN3 (BA3533), purchased from BOSTER; anti-HA (M132-3), purchased from MBL; anti-Flag (AE005), anti-GFP (AE0122), were purchased from ABclonal; anti-β-actin (HC201-01), purchased from TransGen; anti-TRIM25 (67314-1-Ig), purchased from Proteintech; anti-GST (#2622), anti-Phospho-IKKα/β (#2697), anti-Phospho-TBK1 (#5483P), anti-TBK1 (#3504), anti-Phospho-IRF3 (#4947), anti-IRF3 (#4302), anti-Phospho-NF-κB p65 (#3033) and anti-NF-κB p65 (#8242) were purchased from Cell Signaling Technology; goat anti-mouse IRDye680RD (C90710-09) and goat anti-rabbit IRDye800CW (C80925-05), were purchased from Li-COR. Following reagents were used in this study: Polyinosinic–polycytidylic acid potassium salt [poly (I:C)] (P9852, average MW: 200,000 to 500,000), purchased from Sigma-Aldrich; Human recombinant TNF-alpha protein (10602-H01H), purchased from SinoBiological; Lipofectamine 2000, purchased from Invitrogen; anti-GFP beads (KTSM1301), purchased from Alpa-life; Glutathione Sepharose 4B (17-0756-01), purchased from GE Healthcare; The BD™ CBA Mouse Inflammation Kit (552364), purchased from BD™ Bioscience.

### Viral titer and infection

VSV-eGFP and VSV (kindly provided by Dr. Meng Lin, School of Life Science) were propagated and amplified in HEK293T cells and titrated on Vero cells with a standard plaque assay. SeV was a gift from Dr. Yang Du (Zhongshan School of Medicine, Sun Yat-Sen University). For the in vivo study, VSV-eGFP was injected into mice at a titer of 1 × 10^7^ plaque-forming units (PFU)/g for 12 h. For the in vitro study, cells were infected with VSV-eGFP (MOI = 1 for RTN3 induction and immune response activation; MOI = 0.05 for fluorescence and phase contrast (PH) analyses) or SeV (MOI = 0.1 for immune response activation).

### Plasmids and transfection

The plasmids used in the present study were constructed as follows. *RTN3* and *TRIM25* were obtained from the HEK293T cDNA library and cloned into the eukaryotic expression vector pcDNA3.1 in-frame with an HA or Flag tag as well as the eukaryotic expression vectors pEGFP-C2 and pEBG. The HA-tagged *RTN3* fragment was also subcloned into the pMSCV-PURO retrovirus expression vector, and the *RTN3* fragment was truncated based on its structure and subcloned into pcDNA3.1 to generate the HA-tagged mutants T1, T2, T3 and T4. The Flag-RIG-I, Flag-RIG-I CARDs [RIG-I (N)], Flag-TLR3, Flag-TBK1, Flag-P65, Flag-MDA5, Flag-IRF3, HA-tagged Ub, and K63-Ub (K63 only) were kindly provided by Dr. Yang Du. The RIG-I fragment was also subcloned into pEGFP-C2. PrimeSTAR^@^ MAX DNA Polymerase (Takara, RO45A) and a ClonExpress®II One Step Cloning Kit (Vazyme, C11-02) were used to generate all the constructs. For transfection, HEK293T cells were seeded into 24- or 6-well plates overnight and then transfected using Lipofectamine 2000 according to the manufacturer’s protocol. Poly(I:C) was transfected into HEK293T cells using Lipofectamine 2000.

### Generation of gene modified-cell lines

HEK293T cells were seeded into 6-well plates at a density of 0.5 × 10^6^ cells/ml and incubated overnight. Then, 2.5 ml/well of a *TRIM25* shRNA*-*encoding lentivirus was added to the plate, which was then placed in an incubator. Twenty-four hours post inoculation, the cells were infected with a second batch of virus for another 24 h. The infected cells were then reseeded into 6-well plates with fresh RPMI 1640 medium supplemented with 10% FBS and cultivated for 36 h. Puromycin (2-4 μg/ml, Thermo Fisher, A113803) was used for TRIM25-knockdown HEK293T cell selection, and immunoblot analysis was performed to determine the knockdown efficiency. For TRIM25 knockdown-HEK293T cells generation, the following TRIM25 shRNA sequences were used: (1), 5’-CCGGAACAGTTAGTGGATTTA-3’; (2), 5’-GAACTGAACCACAAGCTGATA-3’. Both sequences target the coding region exon1 of TRIM25 gene and cloned into pLKO.1 lentivirus expression vector.

### Luciferase reporter assays

HEK293T cells were seeded in 24-well plates overnight and then transfected using Lipofectamine 2000 with 100 ng of an ISRE-luciferase reporter (firefly luciferase), 100 ng of an IFNB luciferase reporter (firefly luciferase), or 100 ng of an NF-κB luciferase reporter (firefly luciferase) together with 20 ng pRL-TK plasmid (Renilla luciferase), 150 ng Flag-RIG-I (N), Flag-MDA5 or Flag-IRF3-5D mutant and increasing amounts of HA-RTN3. Twenty-four hours post transfection, the cells were collected, and luciferase activity was measured using a Dual-Luciferase Assay kit (Promega, E1960) with a Synergy2 Reader (Bio-Tek) according to the manufacturer’s protocol. The relative level of gene expression was determined by normalizing firefly luciferase activity to Renilla luciferase activity. Poly(I:C) (5 μg/ml) or SeV (MOI = 0.1) was also used as for activation in the assay without RIG-I (N) construct transfection.

### Real-time PCR

Total RNA was extracted from HEK293T or A549 cells dissolved in TRIzol reagent (Invitrogen, 15596026) and then reverse transcribed into cDNA using HiScript® III RT SuperMix (Vazyme). A SYBR Green I Master Mix kit (Roche, 04887352001) was used for real-time PCR with a LightCycler® 480 system (Roche). The following primers were used for the assay: Human RTN3 forward, 5’-AGCTGGCTGTCTTCATGTGG-3’; reverse, 5’-CTCGGGCGATGCCAACATAG-3’. Human beta Actin forward, 5’-CACCATTGGCAATGAGCGGTTC-3’; reverse, 5’-AGGTCTTTGCGGATGTCCACGT-3’. Mouse Rtn3 forward, 5’-AGCTGGCTGTCTTCATGTGG-3’; reverse, 5’-TCCCGGGCAATCCCAACA-3’. Mouse Gapdh forward, 5’-AGGCCGGTGCTGAGTATGTC-3’; reverse, 5’-TGCCTGCTTCACCACCTTCT-3’. Human IFNβ, forward, 5’-CCTACAAAGAAGCAGCAA-3’; reverse 5’-TCCTCAGGGATGTCAAAG-3’. Human IFIT1 forward, 5’-GCCTTGCTGAAGTGTGGAGGAA-3’; reverse, 5’-ATCCAGGCGATAGGCAGAGATC-3’. Human IFIT2 forward, 5’-GGAGCAGATTCTGAGGCTTTGC-3’; reverse, 5’-GGATGAGGCTTCCAGACTCCAA-3’. Human TNFα forward, 5’-CTCTTCTGCCTGCTGCACTTTG-3’; reverse, 5’-ATGGGCTACAGGCTTGTCACTC-3’. Human IL-6 forward, 5’-AGACAGCCACTCACCTCTTCAG-3’; reverse, 5’-TTCTGCCAGTGCCTCTTTGCTG-3’. Human CCL20 forward, 5’-AAGTTGTCTGTGTGCGCAAATCC-3’; reverse, 5’-CCATTCCAGAAAAGCCACAGTTTT-3’. Other primers used in the array were listed in supplemental table 1.

### Coimmunoprecipitation and immunoblot analysis

For coimmunoprecipitation assays, cells transfected and stimulated with the appropriate ligands were lysed with IP buffer (Beyotime, P0013), after which whole cell extracts were collected and incubated with anti-GFP beads or Glutathione Sepharose 4B at 4°C for 1 h or overnight. Then, the beads were washed 4-5 times with IP buffer, and the immunoprecipitants were eluted with 2× SDS loading buffer and then resolved by SDS-PAGE. Subsequently, the proteins were transferred to PVDF membranes (Millipore, ISEQ00010) and further incubated with the indicated primary and secondary antibodies. The images were visualized using an Odyssey Sa system (LI-COR).

### Confocal microcopy

WT HeLa cells were seeded in 8-well Millicell^@^EZ slides (Millipore Ltd, PEZGS0816) overnight and then transfected using Lipofectamine 2000. Twenty-four hours post transfection, the cells were carefully washed with 1× PBS 3 times (2 ml/well, 5 min/time) and then fixed with 4% PFA for 15 min at 4°C. After the cells were penetrated with 0.2% Triton X-100 at room temperature for 5 min, they were washed and blocked with 3% BSA in 1× PBS for 1 h at 4°C and then assayed using the indicated primary and secondary antibodies. Microcopy was performed with a ZESS LSM800 system, and images were captured and processed using ZESS ZEN Imaging Software.

### Histology

Infected mice were sacrificed at the indicated timepoints, and their livers and lungs were dissected and fixed in 4% PFA. After embedding, paraffin sections were stained with hematoxylin and eosin (H&E) or subjected to standard immunohistochemical staining (IHC) before being visualized by bright field microscopy.

### Statistical analyses

The data are presented as the means ± SD (for in vivo experiments, the values are presented as the means ± SD of n mice), and Sidak’s multiple comparisons test was used for all statistical analyses performed with GraphPad Prism 8. Differences between two groups were considered significant at p value < 0.0332.

## Supporting information

All supplemental figures.

## Author contributions

X.Li. and E.Kuang. initiated the concept. Z.Yang., J.Wang., B.He, X.Li. and E.Kuang designed the experiments and analyzed the data. Z.Yang., J.Wang., B.He. and X.Zhang performed the experiments. Z.Yang. and E.Kuang. wrote the paper.

## Acknowledgments

We thank all the members of our laboratory for their critical assistance and helpful discussions. This work is supported by grants from the Natural Science Foundation of China (81671996 and 81871643) to E.K. and the Natural Science Foundation of China (81971928) to X.L.

