## Supplementary material for "RTN3 inhibits RIGI-I-mediated antiviral responses by impairing TRIM25-mediated K63-linked polyubiquitination": All supplemental figures.

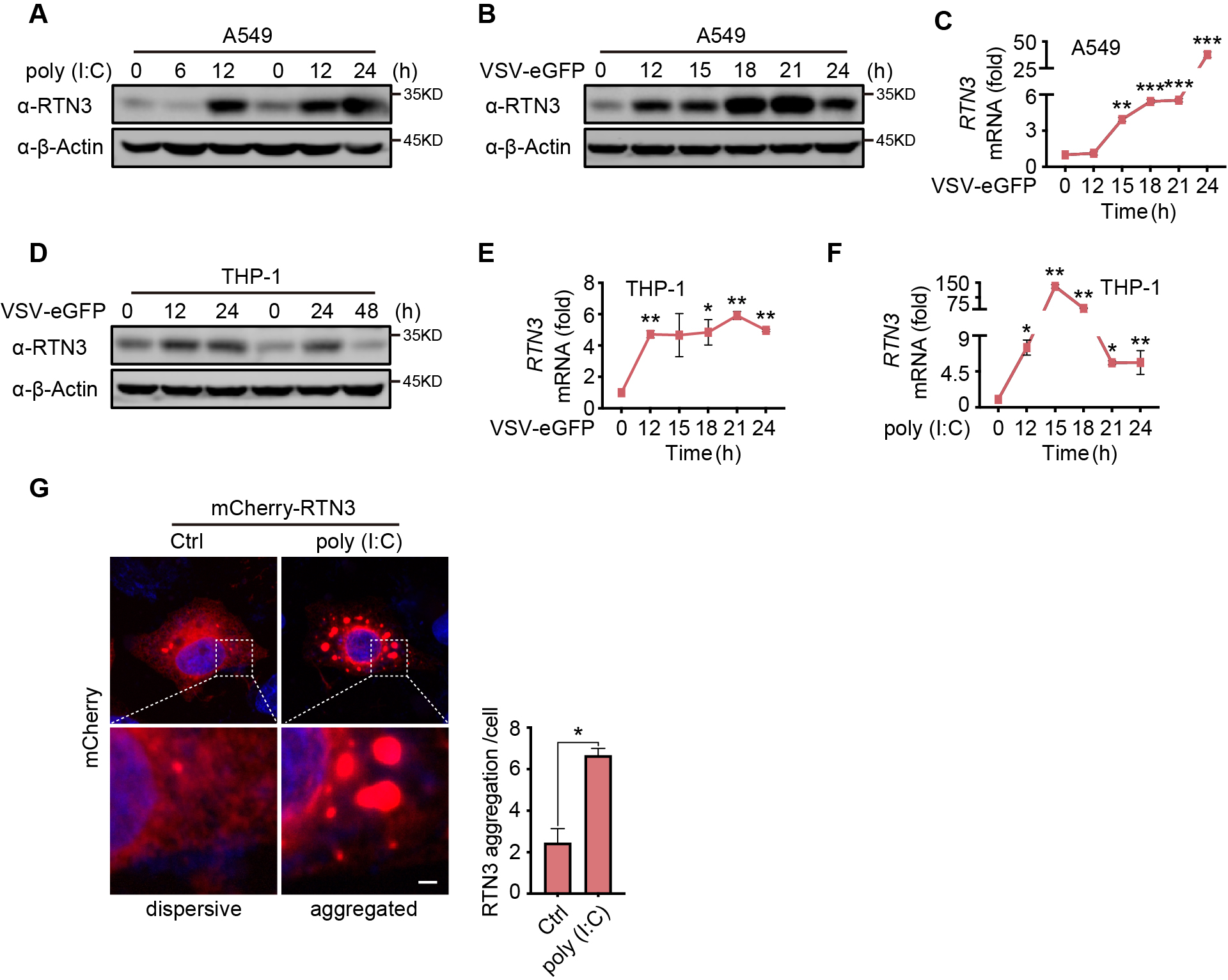

**Figure S1**. **RTN3 is upregulated and self-aggregates upon RNA viral infection.**

A-B. Immunoblot analysis of A549 cells treated with poly(I:C) (5 µg/ml) (A) or infected with VSV-eGFP (MOI = 1) (B) at the indicated timepoints.

C. mRNA levels of *RTN3* in the same samples shown in (B) were detected by RT-PCR.

D. Immunoblot analysis of THP-1 cells infected with VSV-eGFP (MOI = 1) at the indicated timepoints.

E-F. mRNA levels of *RTN3* were detected in THP-1 cells infected with VSV-eGFP (MOI = 1) (E) or treated with poly(I:C) (5 µg/ml) (F) at the indicated timepoints.

G. Confocal microcopy analysis of HeLa cells transfected with mCherry-RTN3 followed by stimulation with poly(I:C). Quantitative comparison of RTN3 aggregation levels as analyzed by aggregation enumeration. For each treatment method, 20 cells/group and 2 groups in total were analyzed for each treatment. Scale bar, 10 μm.

In (A, B, D), the data are representative of three independent experiments. In (C, E, F), the data are shown as the mean values ± SD (n = 3). *, p < 0.0332; **, p < 0.0021; ***, p < 0.0002; and ****, p < 0.0001 by Sidak’s multiple comparisons test. In (H), the data are shown as the mean values ± SD (n = 40); *, p < 0.05 by unpaired t test.

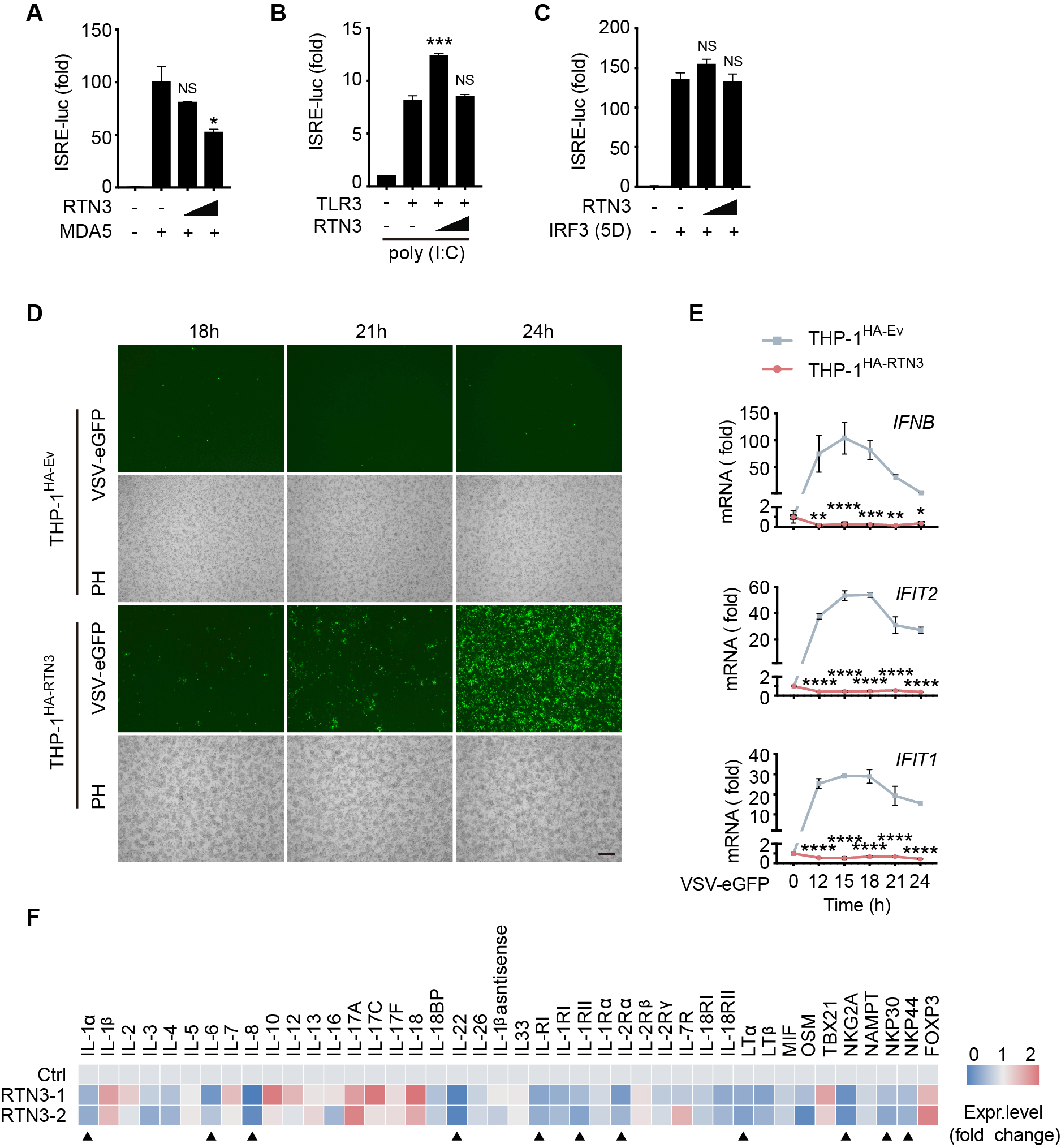

**Figure S2**. **RTN3 exerts and suppresses RIG-I-mediated antiviral immune responses**.

A. Luciferase activity of HEK293T cells transfected with ISRE-Luc together with HA-Ev or increasing amounts of HA-RTN3 (+, ++) and followed by transfection with MDA5 CARDs as an activator.

B. Luciferase activity of HEK293T cells transfected with ISRE-Luc and Flag-TLR3 together with HA-Ev or increasing amounts of HA-RTN3 and followed by treatment with poly(I:C) (5 μg/ml) for 24 h.

C. Luciferase activity of HEK293T cells transfected with ISRE-Luc together with HA-Ev or increasing amounts of HA-RTN3 and followed by transfection of IRF3 (5D mutant) as an activator.

D. Fluorescence and phase contrast (PH) analyses of THP-1 cells stably overexpressing HA-RTN3 (THP-1^HA-RTN3^) and control THP-1 cells (THP-1^HA-Ev^) and followed by infection with VSV-eGFP at the indicated timepoints. Scale bar, 100 μm.

E. RT-PCR detection of *IFN-β*, *IFIT2*, and *IFIT1* mRNA levels in the same samples shown in (D).

F. RT-PCR detection of mRNA levels for the indicated genes in HEK293T cells transfected with HA-Ev or HA-RTN3. “▼” indicates genes that were significantly downregulated in two independent *RTN3-*overexpressing samples. Total RNA was extracted 36 h post transfection.

In (A, B, C, E, F), the data are shown as the mean values ± SD (n = 3). *, p < 0.0332; **, p < 0.0021; ***, p < 0.0002; and ****, p < 0.0001 by Sidak’s multiple comparisons test.

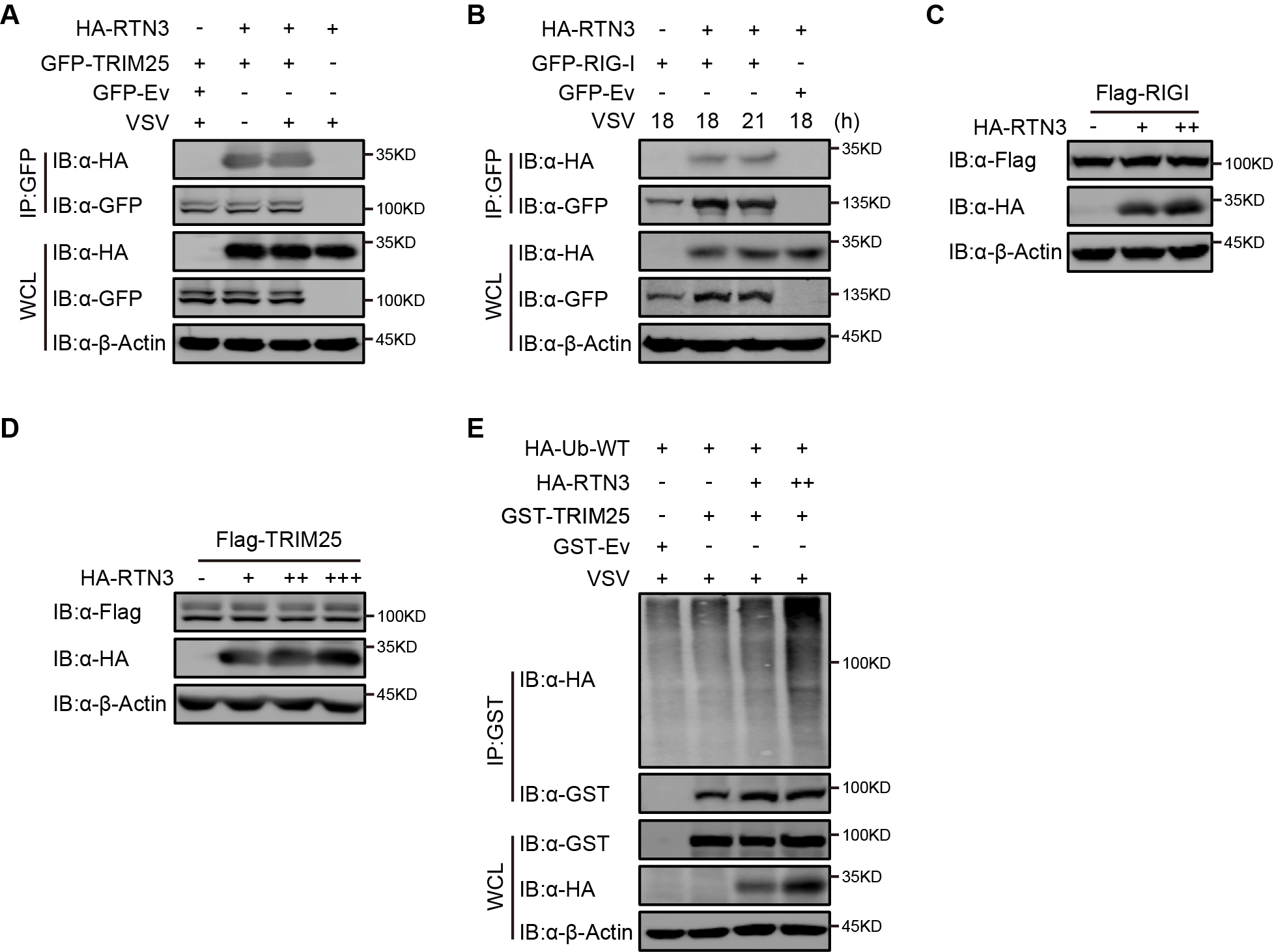

**Figure S3**. **RTN3 has no influence on the protein levels of RIG-I and TRIM25**.

A. CoIP and immunoblot analyses of HEK293T cells transfected with HA-Ev or HA-RTN3 together with GFP tagged TRIM25 (GFP-TRIM25) or GFP-Ev for the indicated groups for 24 h and followed by infection with VSV (MOI = 1) for 8 h.

B. CoIP and immunoblot analyses of HEK293T cells transfected with HA-RTN3 or HA-Ev together with GFP tagged RIG-I (GFP-RIG-I) or GFP-Ev for the indicated groups for 24 h and followed by infection with VSV (MOI = 1) for 8 h.

C. Immunoblot analysis of HEK293T cells transfected with Flag-RIG-I together with HA-Ev or increasing amounts of HA-RTN3 (+, ++).

D. Immunoblot analysis of HEK293T cells transfected with Flag-TRIM25 together with HA-Ev or increasing amounts of HA-RTN3 (+, ++, +++).

E. CoIP and immunoblot analyses of HEK293T cells transfected with HA-Ub-WT and increasing amounts of HA-RTN3 (+, ++) together with GST-TRIM25 or GST-Ev for the indicated groups for 24 h and followed by infection with VSV (MOI = 1) for 8 h.

In (A-E), the data are representative of three independent experiments.

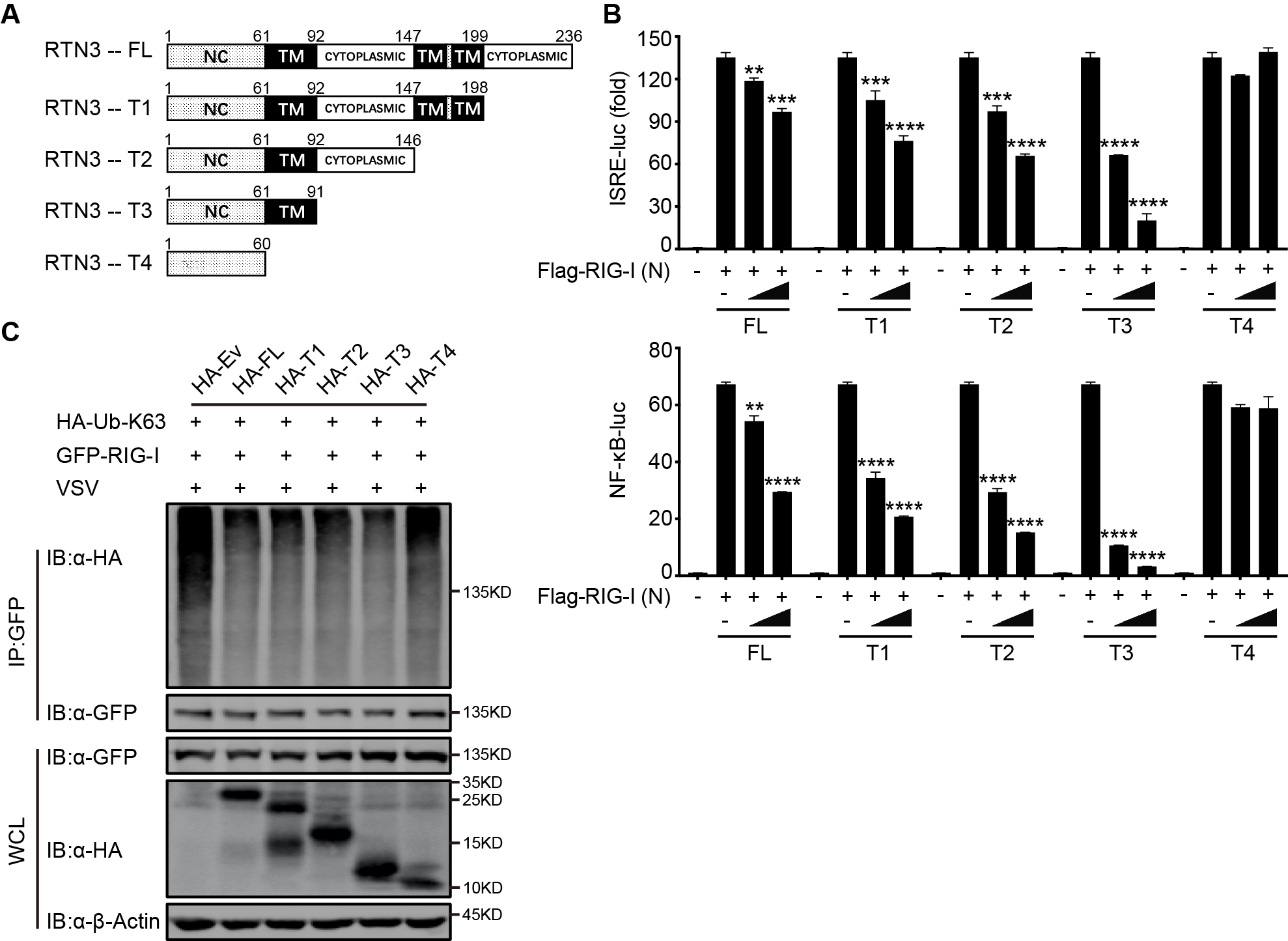

**Figure S4**. **RTN3 inhibitory activity requires its transmembrane domain 1**.

A. The structure of RTN3 and its truncated mutants. Noncytoplasmic domain (NC), transmembrane domain (TM), full length (FL), truncated (T).

B. Luciferase activity of HEK293T cells transfected with ISRE-Luc (top) and NF-κB-Luc (bottom) together with HA-Ev or increasing amounts (wedge) of HA-RTN3 and followed by transfection of RIG-I (N) as an activation of the pathway.

C. CoIP and immunoblot analyses of HEK293T cells transfected with HA-Ub-K63, HA-RTN3 and its truncated mutants as in (A) together with GFP-RIG-I or GFP-Ev for the indicated groups for 24 h and followed by infection with VSV (MOI = 1) for 8 h

In (C), the data are representative of three independent experiments. In (B), the data are shown as the mean values ± SD (n = 3). *, p < 0.0332; **, p < 0.0021; ***, p < 0.0002; and ****, p < 0.0001 by Sidak’s multiple comparisons test.

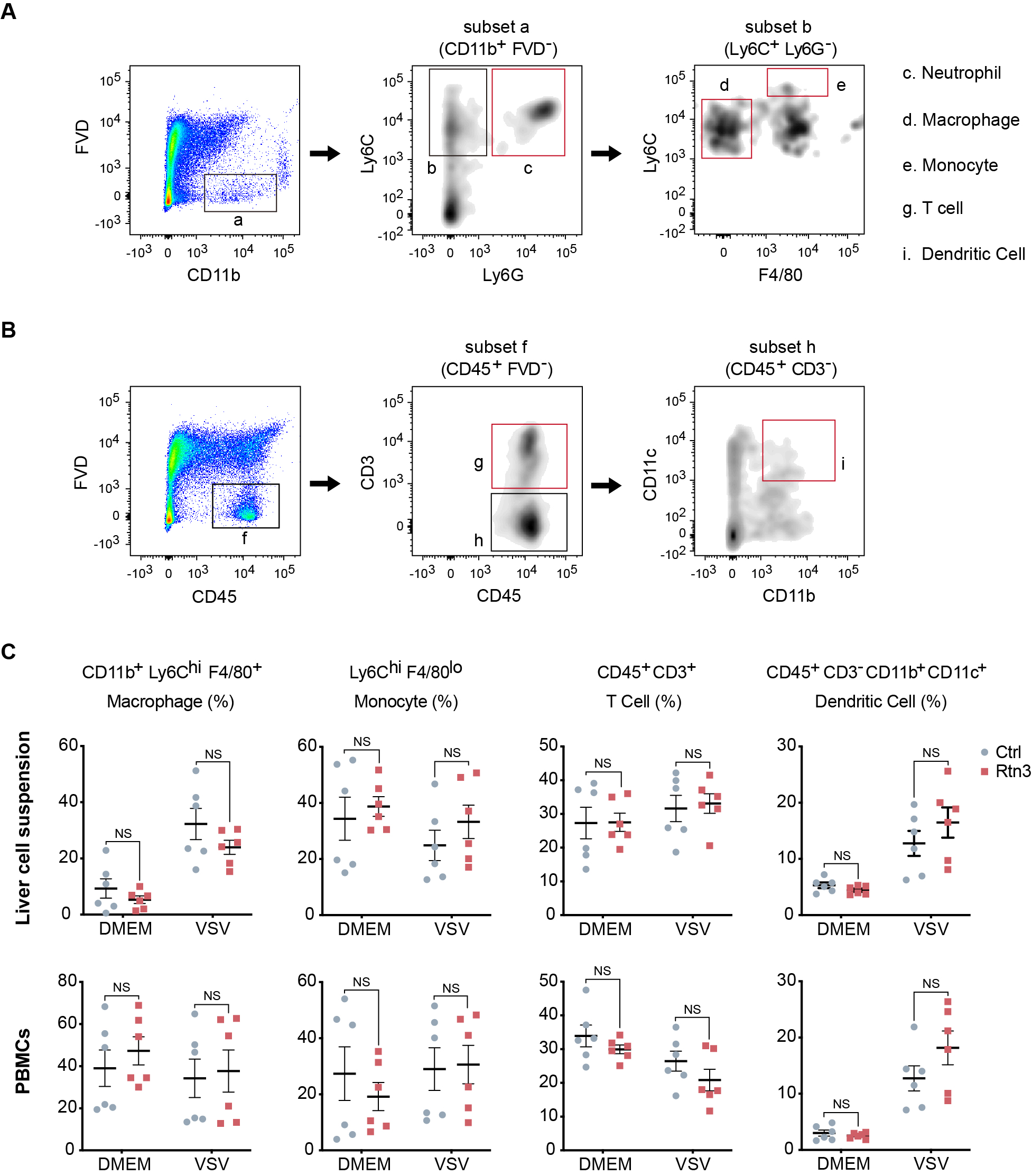

**Figure S5**. **Determination of hydrodynamic injection efficiency and flow cytometry analysis of lymphocytes**.

A-B. Flow cytometry analysis for grouping of liver cell suspension and PBMCs from the same samples shown in (Fig. 5 F, G). Black and red boxes indicate the analyzed populations.

C. Macrophage, monocyte, T cell and dendritic cell populations in the liver cell suspensions and PBMCs from the same samples as shown in (Fig. 5 F, G) were detected by flow cytometry and are presented as a scatterplot.

In (C), the data are shown as the mean values ± SEM (n = 6). *, p < 0.0332; **, p < 0.0021; ***, p < 0.0002; and ****, p < 0.0001 by Sidak’s multiple comparisons test.

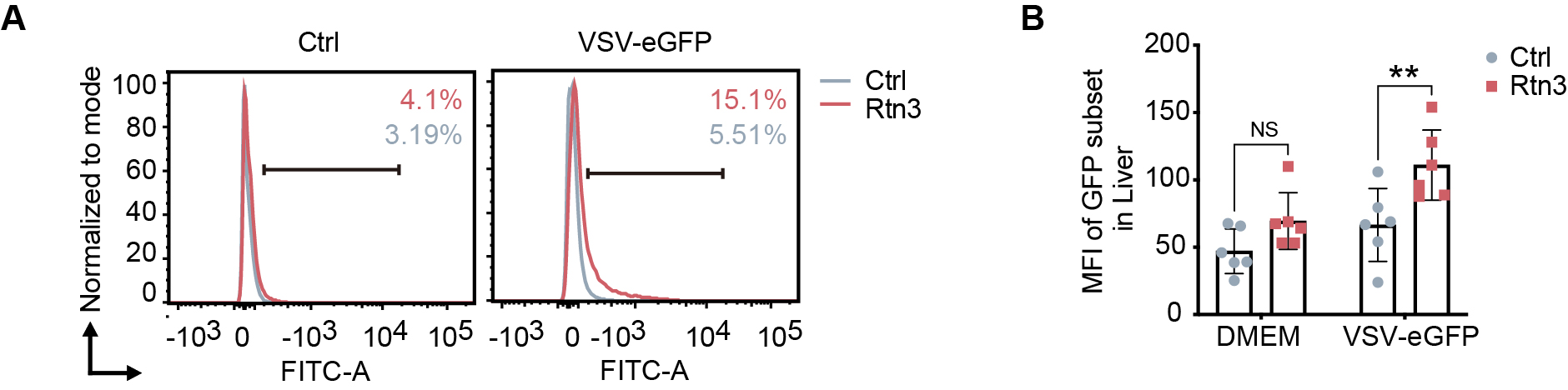

**Figure S6**. **Viral infection in the liver increases upon RTN3 overexpression**.

A-B. The EGFP-positive percentage in the liver cell suspensions and the fluorescent intensity for the same samples shown in Figure 5F and 5G were analyzed by flow cytometry.

In (B), the data are shown as the mean values ± SEM (n = 6). *, p < 0.0332; **, p < 0.0021; ***, p < 0.0002; and ****, p < 0.0001 by Sidak’s multiple comparisons test.

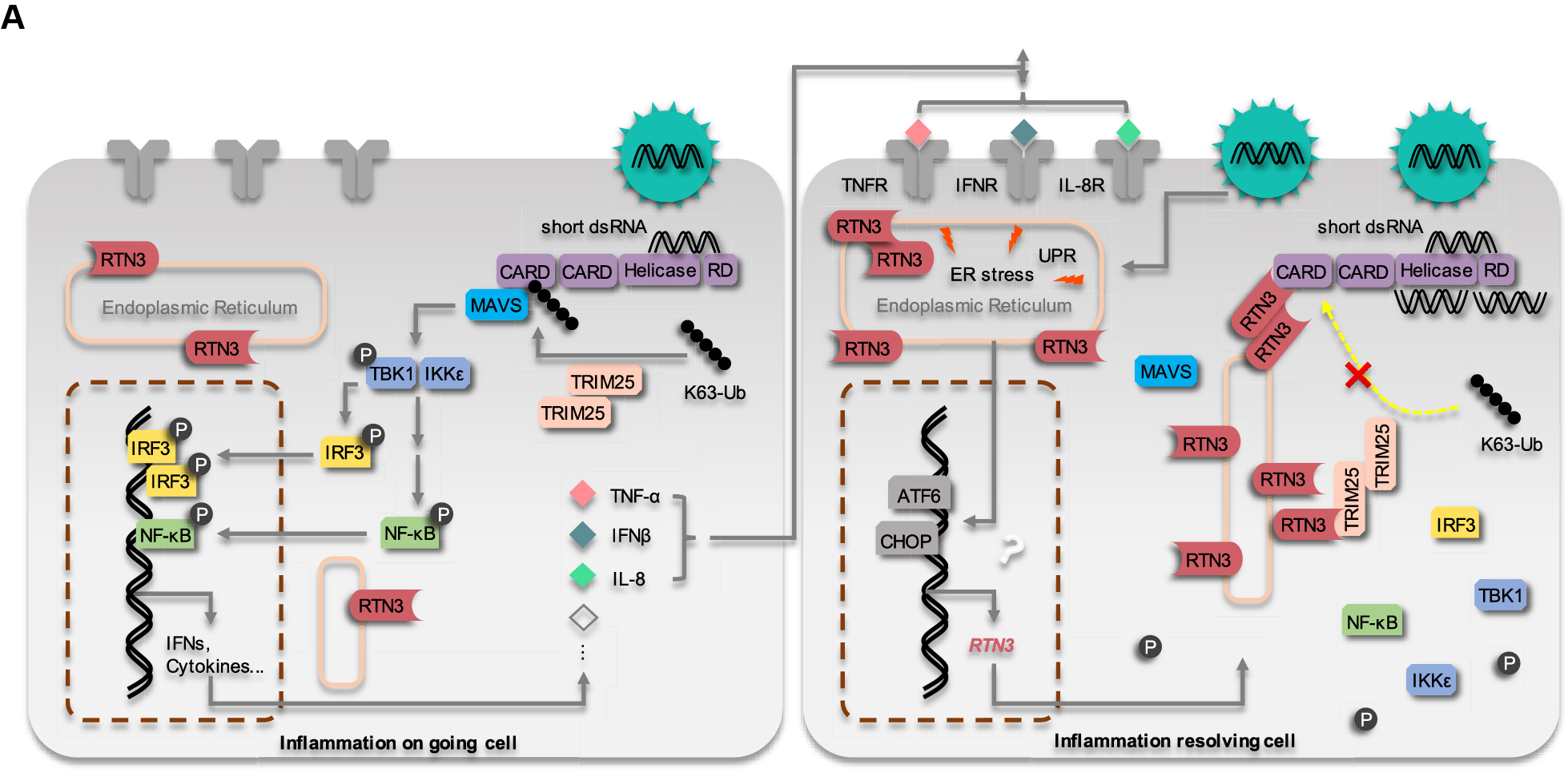

**Figure S6 The working model of RTN3 suppresses RIG-I-mediated antiviral innate immune responses**
